# Efficient callus induction and completion of regeneration in kenaf *Hibiscus cannabinus*

**DOI:** 10.1101/546754

**Authors:** Masaki Odahara, Yoko Horii, Mitsuhiro Kimura, Keiji Numata

## Abstract

Kenaf, *Hibiscus cannabinus*, is a fiber-enriched plant belonging to Malvaceae and is an important fiber crop. The features of kenaf of being fast-growing and fiber-enriched suggest the potential for the use of kenaf in biomass and materials; however, the complete regeneration procedure, which is essential for genetic modification, is lacking. Here, we report the complete regeneration of kenaf from cotyledon explants achieved by callus induction on an improved callus-inducing medium and shoot induction on a shoot-inducing medium and by seed setting under a regulated growth condition in a chamber. Our complete regeneration method will enable the production of stably transformed kenaf that can improve the properties of kenaf as a material.

## Introduction

Kenaf, *Hibiscus cannabinus*, is an annual plant belonging to Malvaceae, which includes cotton and okra. Kenaf grows quickly and has a fiber-enriched woody base; thus, it is regarded as an important fiber crop. The properties of kenaf of being fast-growing and fiber-enriched enable it to be applied to biocomposite materials as a filler, especially in automotive parts[1]. However, establishment of the complete genetic transformation procedure is essential to improve the physical properties of kenaf as a structural material. Among the several ways of genetic transformation established in plants, agrobacterium-mediated transformation is the most popular, and, in general, it utilizes the regeneration of plants from the callus that can be induced by the combination of the plant growth regulators auxins and cytokinins. In kenaf, several studies have reported on the regeneration of shoots. Using the kenaf shoot apex or shoot, shoots are efficiently regenerated without the callus state or adding plant growth regulators, but the addition of N^6^-benzyenine (BA) and thidiazuron, cytokinin-like plant growth regulators, can increase the number of regenerating shoots[2–4]. McLean, Lawrence (5) showed efficient induction of the callus from kenaf internodal stems by adding 1-naphthaleneacetic acid/BA or 2,4-dichlorophenoxyacetic/kinetin acid as the auxin/cytokinin combination, but the callus regenerates shoots with less than 15% efficiency. Samanthi, Puad (6) showed the induction of the kenaf callus from an intact cotyledon with maximally 80% efficiency by adding indole-3-butyric acid (IBA) as the auxin and BA, and the callus regenerated shoots with 68.7% efficiency. Despite sufficient information regarding callus induction and shoot regeneration, which still leaves room for improvement, information regarding the flowering and seed setting is lacking in kenaf. Here, we report the complete regeneration of kenaf including improved callus induction and shoot regeneration, flowering, and seed setting methods.

## Results and Discussion

### Callus induction from kenaf tissues

The callus induction efficiency depends on the tissues type. We tested the seeds, hypocotyls, and cotyledons of kenaf seedlings for their capability to induce callus by cultivating them on a callus-inducing medium (CIM), a widely used auxin-rich medium to induce the plant callus. By using some combinations of auxins and cytokinins reported to induce callus from kenaf explants[5, 6], all the tested tissues induced the callus efficiently (Fig. 1). However, the calli from the tissues showed some differences in properties.

**Figure 1.**
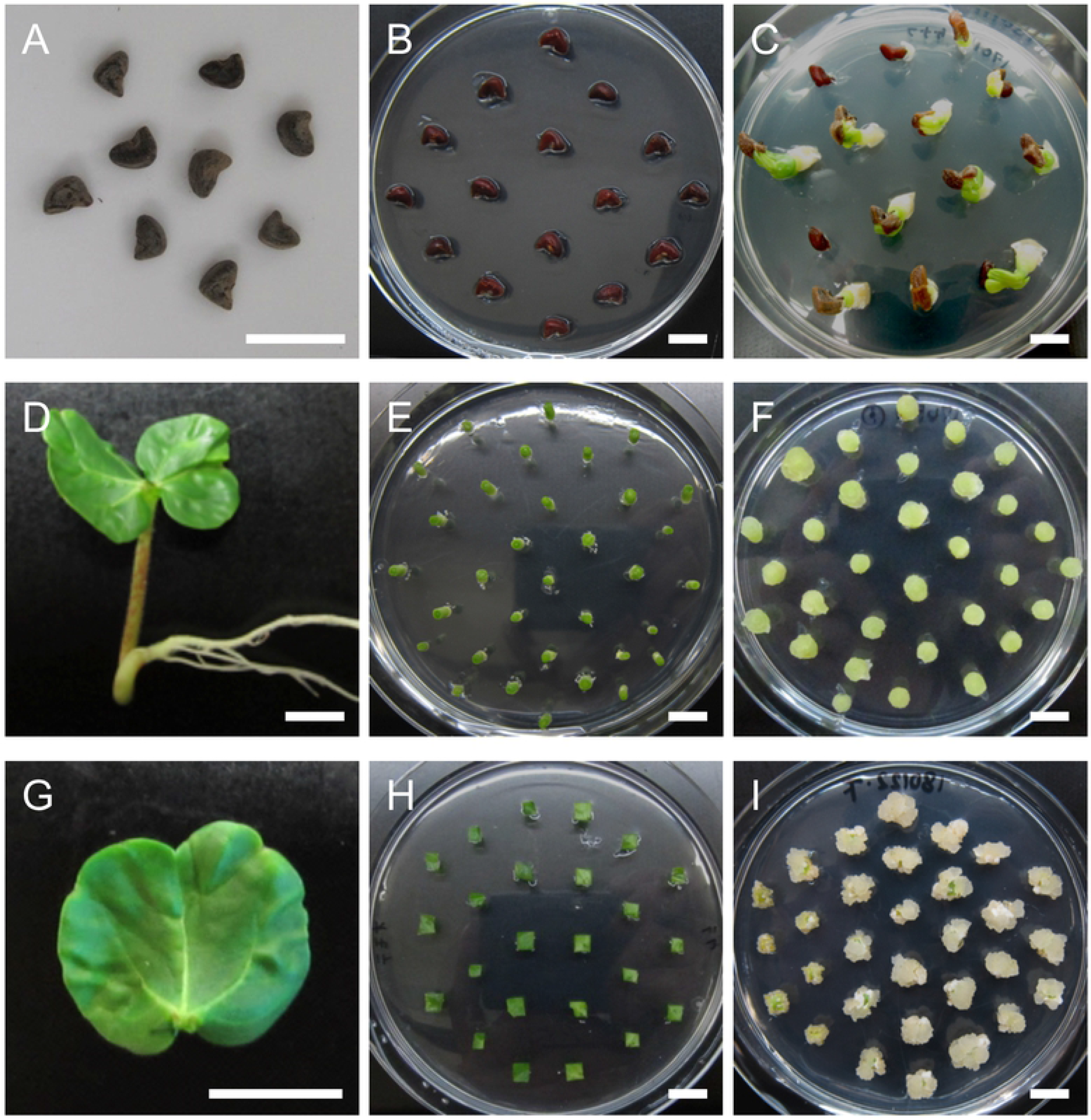
Callus induction from kenaf seed, hypocotyl, and cotyledon. (A-C) Callus induction from kenaf seeds. Kenaf seeds (A) were sterilized and then were placed on CIM (B). Calli induced from seeds cultivated for one week. (D-F) Callus induction from the kenaf hypocotyl. Hypocotyls from 10-day-old seedlings (D) were cut and placed vertically on CIM (E). Calli were induced from the fragmented hypocotyls cultivated for one week (F). (G-I) Callus induction from the kenaf cotyledon. Cotyledons from 10-day-old seedlings (G) were cut into small pieces and then were cultivated on CIM (H). Calli were induced from three-week cultivated cotyledon pieces (I). Bars = 10 mm.

In the case of seeds, after root development in germination, calli were induced from the roots, but subsequent cultivation of the calli on a shoot-inducing medium (SIM), a cytokinin-rich medium for the induction of shoot regeneration, induced the regeneration of only roots but not shoots (Fig. S1). This suggests that the calli induced from roots were not suitable for shoot regeneration, inconsistent with the related species *Hibiscus sabdariffa*[7]. Hypocotyls were fragmented and then were placed vertically or horizontally, or placed horizontally after longitudinal split, or were chopped, and then they were cultivated on CIM. These sectioning and cultivating procedures, except for chopping, succeeded in inducing the callus, and the vertically cultivated hypocotyls seemed to show the highest efficiency for induction (Fig. 1B and Fig. S2). Too much fragmentation may disturb callus induction from hypocotyls. The calli from the vertically cultivated hypocotyls were then tested for the induction of shoot regeneration on SIM with several combinations of plant growth regulators. No shoots were regenerated from these hypocotyl-derived calli (Fig. S3), suggesting the calli derived from hypocotyls were not suitable for shoot regeneration, as well as the calli from roots. Cotyledons showed efficient callus induction on CIM supplemented with indole-3-butyric acid (IBA), a type of auxin, in combination with BA when they were cut into 5-mm pieces, and subsequent cultivation on SIM induced greening of the calli (Fig. 4E), which should lead to shoot development. These results show that kenaf cotyledons, but not seeds or hypocotyls, efficiently induce calli that potentially regenerate shoots.

### Optimization of conditions for callus induction from cotyledons

To increase the efficiency of callus induction from cotyledons, we carried out optimization of plant growth regulator concentrations, basal salts, and solidification reagents for callus induction. First, we analyzed the effect of several concentrations of IBA and BA as supplements to Murashige-Skoog (MS) medium[8]. The results showed that 1.5 mgL^−1^ of BA with 0.01 mgL^−1^ of IBA and 1.5 mgL^−1^ of BA only yielded the best and worst callus induction from the kenaf cotyledon in terms of efficiency and homogeneity of the calli with low induction of root formation, respectively (Table 1 and Fig. S4). This shows that IBA is important for the efficient induction of callus from the cotyledon. However, an excess amount of IBA inhibits callus formation. The root formation was induced by a low cytokinin/auxin ratio in agreement with a report by Skoog and Miller (9). It was noted that the callus-induction efficiency reached 98.0% in 1.5 mgL^−1^ of BA with 0.01 mgL^−1^ of IBA (Table 1), showing an almost complete induction of callus from kenaf cotyledon pieces.

**Table 1.** Callus induction and root formation efficiency from kenaf cotyledons.

|  | IBA concentration (mgL<sup>-1</sup>) combination with 1.5 mg L<sup>-1</sup> of BA |  |  |  |
| --- | --- | --- | --- | --- |
|  | 0 | 0.005 | 0.01 | 0.05 |
| Rate of callus formation | 61.1% | 98.8% | 98.0% | 97.0% |
| (callus/explant) | (214/350) | (395/400) | (392/400) | (388/400) |
| Rate of root formation (%) | 0% | 10.1% | 4.8% | 38.1% |
| (with root/callus) | (0/214) | (40/395) | (19/392) | (148/388) |

Next, we tested the effect of inorganic basal salts, Gamborg B5 [10], Murashige-Skoog (MS)[8], or 1/2 MS, supplemented with 1.5 mgL^−1^ of BA with 0.01 mgL^−1^ of IBA for the induction of callus from cotyledon pieces. In contrast to the callus hardly induced in the B5 medium, the callus was efficiently induced in either MS medium, especially in the MS medium (Fig. 2), suggesting that MS is the more appropriate basal salts than Gamborg B5 and that callus induction from the cotyledon requires a high concentration of nutrients. Using the MS medium supplemented with 1.5 mgL^−1^ of BA with 0.01 mgL^−1^ of IBA, our analysis also clarified that the use of agar instead of phytagel (gellan gum) as a solidification reagent significantly disturbed callus induction from the kenaf cotyledon (Fig. 3), suggesting that solidification reagents are important for kenaf callus induction as well as the effect of plant growth regulators.

### Shoot regeneration from the callus

Using the induced kenaf calli, we next tested the regeneration of shoots with several patterns of plant growth regulator concentrations. Calli formed on CIM were transferred to SIM supplemented with 0.3 mgL^−1^ of gibberellin (GA) in which the concentration of BA and IBA were modified to 0.5-2 mgL^−1^ and 0-0.05 mgL^−1^, respectively. As a result of cultivation on the SIM, greening of the calli was efficiently induced in the SIMs containing both 1-2 mgL^−1^ of BA and 0.05 mgL^−1^ of IBA, but not in other SIMs (Fig. S5 and Table S2). This suggests a lower concentration of BA (<1 mgL^−1^) or IBA (<0.05 mgL^−1^) affects the greening of calli. Subsequent cultivation of these greened calli on the SIM supplemented with 1.5 mgL^−1^ of BA, 0.05 mgL^−1^ of IBA, and 0.3 mgL^−1^ of GA led to the regeneration of multiple shoots from a single callus (Fig. 4C). To induce the development of the regenerated shoot with roots, they were transferred to an MS medium without any plant growth regulators. Cultivation on the MS led to the induction of shoot elongation and root development (Fig. 4D and E) that could grow and mature in the soil (Fig. 4F). Thus, our data showed that SIM supplemented with 1.5 mgL^−1^ of BA, 0.05 mgL^−1^ of IBA, and 0.3 mgL^−1^ of GA is sufficient for the greening and shoot development of the kenaf callus.

### Flowering and seed maturation of kenaf

Producing the next generation is essential to obtain stable homozygous transgenic plants. However, the regulated condition for flowering and seed maturation of kenaf, especially the regenerated one, is not well defined. We cultivated regenerated kenaf plants on soil at 22 °C or 30 °C under an 8-hour daily photoperiod in a chamber. Cultivation of kenaf at 22 °C succeeded in the development of flower buds, while cultivation at 30 °C failed, suggesting a high temperature affected the development of the flower bud (Fig. 5B). The kenaf plants bearing flower buds were then cultivated at 22 °C or 30°C because, in general, the accumulated temperature after flowering influences the seed maturation; cultivating at a higher temperature is expected to accelerate seed maturation. Both kenaf plants cultivated at 22 °C and 30 °C induced flowering normally (Fig. 5C), and subsequent cultivation at 22 °C led to the development of fruits from the flowers and maturation of seeds in the fruits (Fig. 5D), although the initial few fruits were dropped without seed maturation. By contrast, cultivation at 30 °C after flowering failed to set fruit because of the development of an abscission layer at the base of the fruit in the initial stage of fruit development. These results collectively suggest that high temperature inhibits the development of flower and fruit buds but not the growth of the flower. The regenerated kenaf fruits matured at 22 °C contained 8 to 14 seeds (Fig. 5E). The seeds from the regenerated kenaf were evaluated by sawing and cultivating them on MS medium. Forty-seven seeds from four fruits of the regenerated kenaf were germinated at a rate of 62%, suggesting a sufficient germination rate of the seeds from the regenerated kenaf. In conclusion, cultivation at 22 °C under an 8-h daily photoperiod fulfills the conditions for both flowering and seed development of kenaf.

### Conclusion

In this report, we discussed the successful complete regeneration of kenaf, *Hibiscus sabdariffa*, via the callus state induced from cotyledon explants. Multiple shoot regeneration from the greened callus, flowering, and seed maturation of the regenerated plant in a regulated condition enabled production of the next generation. In addition to the agrobacterium infection method that is still incomplete for the production of the kenaf T_2_ generation[11–13], a new gene transformation method with functional peptide has recently been developed in plants[14–16]. In combination with the transformation methods, our complete regeneration method would produce stable genetically transformed kenaf. Since kenaf can be used as a filler of composite materials[1, 17, 18], modification of the physical property of kenaf by stable transformation is expected to be applied to the development of improved materials.

## Materials and Methods

### Plant material and seed germination

Kenaf *Hibiscus cannabinus* seeds were purchased from Fujita seed corporation (Hyogo, Japan). Kenaf seeds were sterilized with 70% ethanol for 20 min and then with 20% sodium hypochlorite for 30 min, followed by rinsing with sterilized water. The seeds were placed on MS medium containing 3% sucrose with or without growth regulators and were cultivated at 22 °C under a 16-hour daily photoperiod.

### Culture media and condition for callus induction

For callus induction from seeds, sterilized seeds were placed on MS medium containing 3% sucrose, 0.3 mgL^−1^ of 2,4-D, 0.3 mgL^−1^ of Kinetin and 0.3% of phytagel, and then were cultivated for one week at 22 °C under a 16-h daily photoperiod. Regarding hypocotyls, those from aseptically grown 10-day-old seedlings were cut into a 5-mm length or chopped into smaller fragments, and then were placed on MS medium containing 3% sucrose, 2.0 mgL^−1^ of 2,4-D, 2.0 mgL^−1^ of Kinetin and 0.3% phytagel. They were cultivated for one to four weeks at 22 °C under a 16-h daily photoperiod. Regarding cotyledons, those from aseptically grown 10-day-old seedlings were cut into 5-mm pieces and were placed on MS, 1/2 MS, or Gamborg B5 medium containing 2% sucrose, supplemented with 1.5 mgL^−1^ of BA and 0-0.05 mgL^−1^ of IBA and 0.3% phytagel or 0.8% agar. Calli were induced for three weeks at 22 °C under dark conditions.

### Culture media and conditions for shoot induction

Calli induced from seeds were transferred to MS medium containing 2% sucrose, 1.0 mgL^−1^ of BA and 0.3% phytagel, and then were cultivated for four weeks at 30°C under continuous light. Calli induced from hypocotyls were transferred to MS medium containing 2% sucrose, 0.1-5.0 mgL^−1^ of BA, 0.1-5.0 mgL^−1^ of CPPU or 1.0 mgL^−1^ of trans-Zeatin, 0-0.3 mgL^−1^ of IAA, and 0.3% phytagel, and then were cultivated for four weeks at 30 °C under continuous light. Calli induced from cotyledons were transferred to MS medium containing 2% sucrose, 0.5–2.0 mgL^−1^ of BA, 0–0.05 mgL^−1^ of IBA, 0.3 mgL^−1^ of GA, and 0.3% phytagel, and then were cultivated for four weeks at 30 °C under continuous light.

### Flowering and seed maturation

Shoots regenerated from the callus were cultivated on MS medium containing 2% sucrose and 0.3% phytagel at 30 °C under continuous light. The plants were then transferred to a 2:1 mixture of soil and vermiculite and were cultivated at 22 °C or 30 °C under continuous light. The plants were then cultivated at 22 °C or 30 °C under an 8-hour daily photoperiod.

## Acknowledgements

This work was supported by the Japan Science and Technology Agency Exploratory Research for Advanced Technology (JST-ERATO; Grant No., JPMJER1602).

## Figure legends

**Figure 2. Effect of inorganic basal salts for callus induction from cotyledons.** Calli were induced from cotyledon pieces on CIM composed of Gamborg B5 (A), 1/2 MS (B), or MS (C) supplemented with 1.5 mgL^−1^ of BA and 0.01 mgL^−1^ of IBA for three weeks. Bars = 10 mm.

**Figure 3. Effect of solidification reagents on callus induction from cotyledons.** Calli were induced from cotyledon pieces on MS medium containing 1.5 mgL^−1^ of BA and 0.01 mgL^−1^ of IBA, and 0.3% phytagel (A) or 0.8% agar (B) for three weeks. Bars = 10 mm.

**Figure 4. Regeneration of kenaf from cotyledons via calli.** Calli induced from cotyledon pieces on CIM (A) were then cultivated on SIM (B). After the greening of calli (C), multiple shoots were regenerated (D). The shoots were then cultivated on growth regulator-free MS medium (E) and subsequently cultivated on soil (F). Bars = 10 mm.

**Figure 5. Flowering and seed maturation of regenerated kenaf.** (A) Regenerated kenaf grown on soil in a growth chamber. (B-E) Representative images of flower buds (B), a flower (C), an immature seed pod, and matured seeds (E). Bars = 10 cm in A and 1 cm in B-E.

